# Sex Differences in Functional Topography of Association Networks

**DOI:** 10.1101/2021.05.25.445671

**Authors:** Sheila Shanmugan, Jakob Seidlitz, Zaixu Cui, Azeez Adebimpe, Danielle S. Bassett, Maxwell A. Bertolero, Christos Davatzikos, Damien A. Fair, Raquel E. Gur, Ruben C. Gur, Bart Larsen, Hongming Li, Adam Pines, Armin Raznahan, David R. Roalf, Russell T. Shinohara, Jacob Vogel, Daniel H. Wolf, Yong Fan, Aaron Alexander-Bloch, Theodore D. Satterthwaite

## Abstract

Prior work has shown that there is substantial interindividual variation in the spatial distribution of functional networks across the cerebral cortex, or *functional topography*. However, it remains unknown whether there are sex differences in the topography of individualized networks in youth. Here we leveraged an advanced machine learning method (sparsity-regularized nonnegative matrix factorization) to define individualized functional networks in 693 youth (ages 8-23 years) who underwent functional magnetic resonance imaging as part of the Philadelphia Neurodevelopmental Cohort. Multivariate pattern analysis using support vector machines classified participant sex based on functional topography with 83% accuracy (*p*<0.0001). Brain regions most effective in classifying participant sex belonged to association networks, including the ventral attention and default mode networks. Mass-univariate analyses using generalized additive models with penalized splines provided convergent results. Comparative analysis using transcriptomic data from the Allen Human Brain Atlas revealed that sex differences in multivariate patterns of functional topography correlated with the expression of genes on the X-chromosome. These results identify normative developmental sex differences in the functional topography of association networks and highlight the role of sex as a biological variable in shaping brain development in youth.

**SIGNIFICANCE STATEMENT:** We identify normative developmental sex differences in the functional topography of personalized association networks including the ventral attention network and default mode network. Furthermore, chromosomal enrichment analyses revealed that sex differences in multivariate patterns of functional topography were spatially coupled to the expression of X-linked genes as well as astrocytic and excitatory neuronal cell-type signatures. These results highlight the role of sex as a biological variable in shaping functional brain development in youth.

## INTRODUCTION

Significant sex differences have been documented in cognitive domains including visuospatial processing, social cognition, emotional memory, and executive function (1–3). Prior studies have sought to understand these behavioral differences in the context of sex differences in brain structure, organization, and function that have been noted during childhood and adolescence (1, 4, 5). Understanding normative sex differences in brain structure and function not only allows us to learn more about the neurobiology of developmental sex differences in cognition and behavior, but is also a necessary first step in constructing a framework to study sex differences in psychopathology, where such differences are even more prominent and emerge during development.

Previous neuroimaging studies have examined sex differences in network connectivity as a contributor to sex differences in cognition and psychiatric disorders (6–8). For example, males exhibit greater between-module connectivity and lower within-module connectivity than females (9, 10). These patterns of connectivity have been linked to better performance on spatial and motor tasks, cognitive domains where males outperform females (9). Furthermore, depression and anxiety are more prevalent in females (11), and brain connectivity patterns associated with mood disorder symptoms are greater in females (12). In contrast, ADHD (13) and conduct disorder (14) are more prevalent in male youth, and may be related to abnormal functional connectivity within and emerging from executive regions (15). Although these studies and others suggest that sex differences in network connectivity may underlie various behavioral phenotypes, findings have been heterogeneous, raising concerns about reproducibility and potential for clinical translation.

One potential reason for such heterogeneity in findings among prior studies is the use of standardized network atlases. Standardized atlases assume a stable 1:1 correspondence between structural and functional anatomy across individuals. Such methods assume that by aligning brain structural anatomy across subjects, functional network anatomy across subjects is also brought into alignment. However, evidence from multiple independent groups has shown that there is significant inter-individual variation in the spatial distribution of functional networks across the anatomic cortex, or *functional topography* (16–20). These studies demonstrate that mapping between structure and function varies substantially between individuals, and that interindividual variation of personalized functional networks is maximal in association networks such as the ventral attention, frontoparietal, and default mode networks (20). Assuming a 1:1 correspondence between structural and functional anatomy can alias individual differences in topography into measurement of inter-regional functional connectivity. In light of these difficulties, it remains unknown if sex differences in functional topography exist. Furthermore, it is unknown whether such differences might emerge in youth – a period marked by extensive remodeling of functional networks (20).

Accordingly, here we capitalized upon a large sample of youths imaged as part of the Philadelphia Neurodevelopmental Cohort (21) to evaluate developmental sex differences in functional topography. We used machine learning to define individualized functional networks, hypothesizing that sex differences would be greatest in association networks. Because sex differences in neuroanatomy have previously been linked to sex chromosome gene expression (22, 23), we also evaluated the relationship between sex differences in topography and gene expression. We predicted that cortical patterns of prominent sex differences in functional network topography would be spatially similar to the expression of sex-chromosome genes.

## RESULTS

As previously described (18), we used sparsity-regularized non-negative matrix factorization (NMF) (24) to derive individualized functional networks in 693 youth (57% female) ages 8-23 years imaged with fMRI as part of the Philadelphia Neurodevelopmental Cohort. In this procedure for defining individualized networks, we first create a consensus atlas for the group of 693 subjects and then use this consensus atlas to define individualized networks for each participant under a sparsity constraint (**Figure 1a**). Using a consensus atlas ensures spatial correspondence across individuals. Seventeen functional networks were identified for each participant (**Figure 1b**), which correspond with commonly used atlases and prior work (17, 20, 25, 26). Networks were named as in Cui *et al*. (20), and include default mode networks (1, 8, and 12), frontoparietal networks (3, 15, and 17), ventral attention networks (7 and 9), dorsal attention networks (5 and 14), visual networks (6 and 10), somatomotor networks (2, 4, 11 and 13), and an auditory network (16). In contrast to hard partitioning methods that assign each vertex to a single network, NMF is a soft partitioning method that yields a probabilistic parcellation such that there are 17 loadings for each vertex that quantify the extent to which it belongs to each network. This probabilistic parcellation can be converted into discrete network definitions for display by labeling each vertex according to its highest loading (**Figure 1c**). Visual examination of individual participants’ functional networks revealed distinct differences in topographic features (**Figure 2**). This inter-individual variation in topography was particularly apparent in association networks such as the ventral attention and default mode networks. In contrast, motor and sensory networks appeared to be much more consistent across individuals.

**Figure 1.**
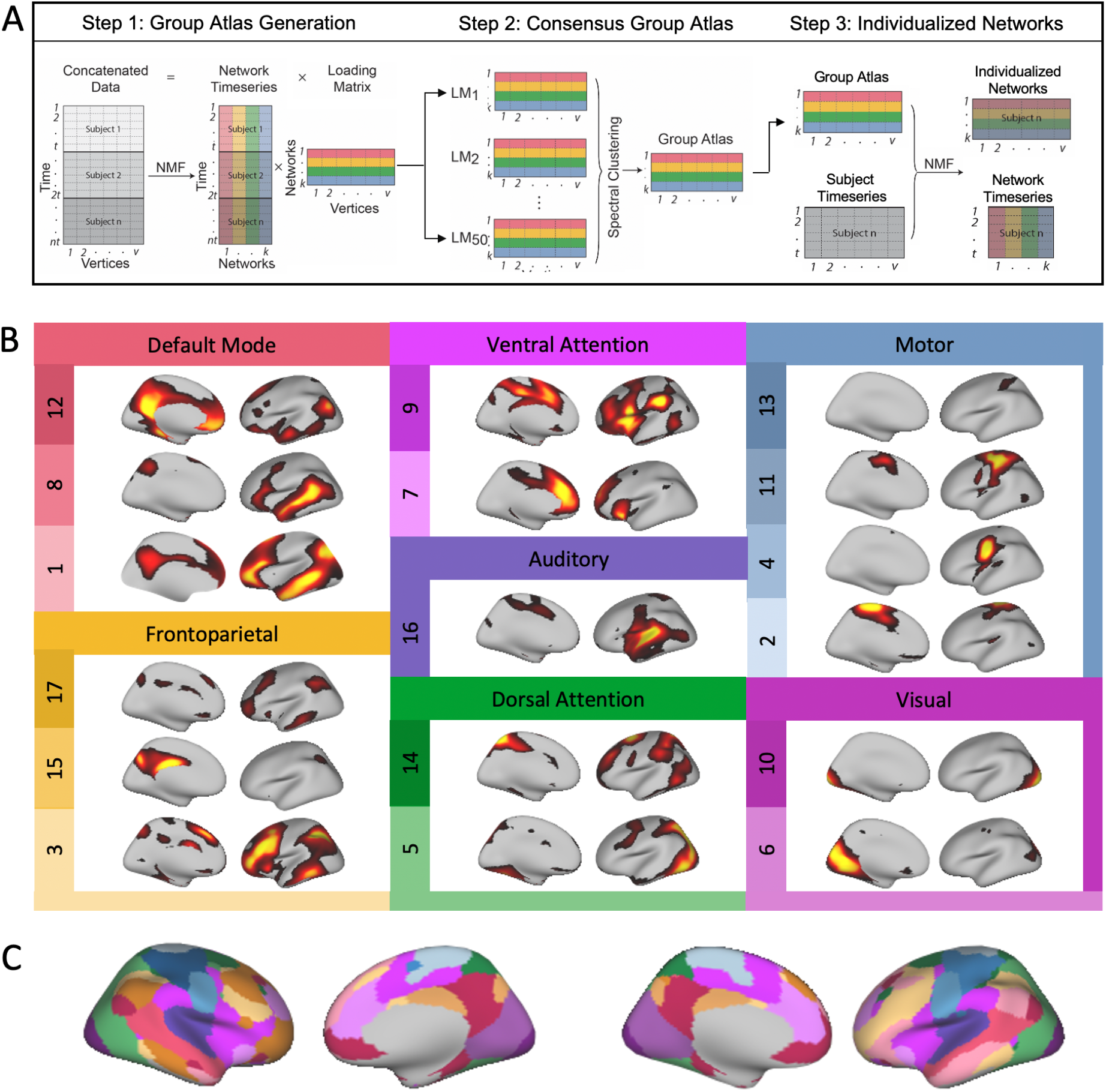
Defining personalized functional networks with non-negative matrix factorization. **A)** We used spatially regularized non-negative matrix factorization (NMF) to derive individualized functional networks. Three fMRI runs were concatenated for each subject, resulting in a 27.4 minute time series with 555 time points for each subject. In step 1, time series from 100 randomly selected subjects were concatenated into a matrix with 55,500 time points (rows) and 17,734 vertices (columns). NMF was used to decompose this data into a time series matrix and loading matrix. The loading matrix had 17 rows and 17,734 columns, which encoded the membership of each vertex for each network. This procedure was repeated 50 times, with each run including a different subset of 100 subjects. In step 2, a normalized-cut based spectral clustering method was applied to cluster the 50 loading matrices into one consensus loading matrix, which served as the group atlas and ensured correspondence across individuals. In step 3, NMF was used to calculate individualized networks for each participant, with the group atlas used as a prior. **B)** Seventeen functional networks were identified for each participant. Networks identified included default mode networks (1,8, and 12), frontoparietal networks (3, 15, and 17), ventral attention networks (7 and 9), dorsal attention networks (5 and 14), visual networks (6 and 10), somatomotor networks (2, 4, 11 and 13), and an auditory network (16). NMF yields a probabilistic (soft) parcellation such that there are 17 loadings for each vertex that quantify the extent to which it belongs to each network. For each loading map, brighter colors indicate greater loadings. **C)** The probabilistic parcellation can be converted into discrete (hard) network definitions for display by labeling each vertex according to its highest loading.

**Figure 2.**
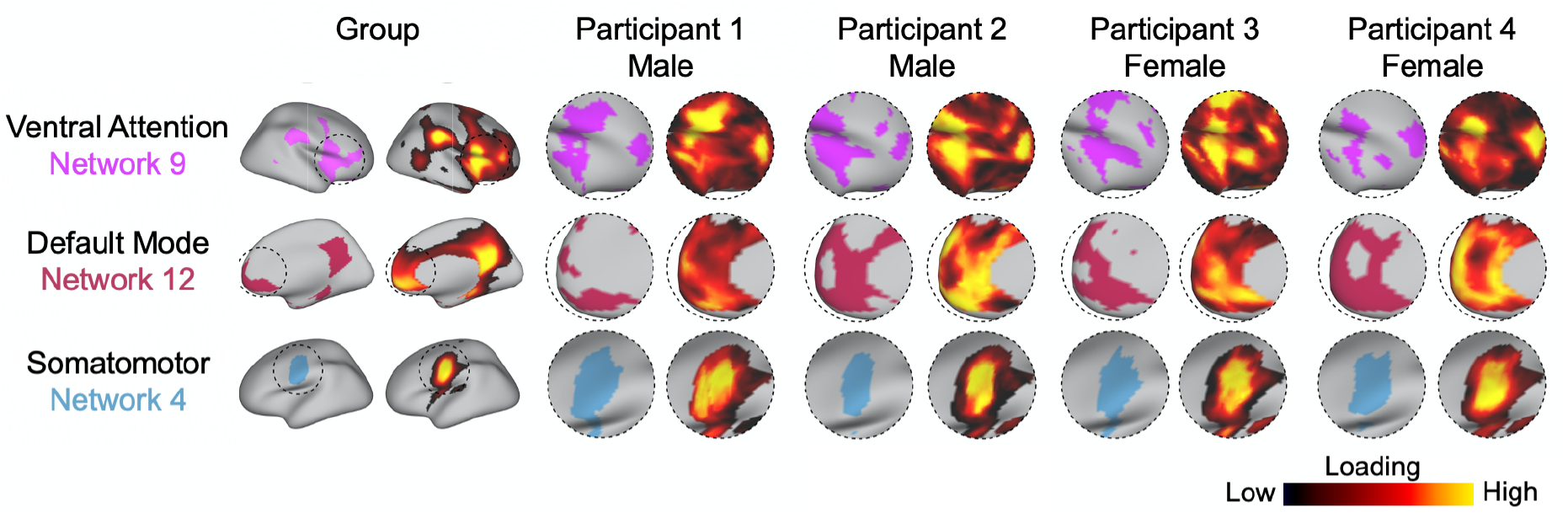
Functional network topography varies between individuals and by sex. Probabilistic loading map and discrete network parcellations of three networks are displayed for the group and four randomly selected participants. Visual examination of individual participants’ functional networks reveal distinct differences in topographic features. This inter-individual variation in topography is particularly apparent in association networks such as the ventral attention and default mode networks. In contrast, motor and sensory networks appear to be more consistent across individuals.

### Machine learning accurately identifies sex using functional topography

Based on our observation that the spatial distribution of association networks varies across individuals, we hypothesized that sex contributed to this inter-individual variation in topography. To test this hypothesis, we first sought to understand the way in which high dimensional patterns of functional topography reflect sex. Multivariate pattern analysis allows for such integration of high dimensional data and can also identify complex patterns of topography that discriminate between males and females. We therefore used a linear support vector machine (SVM; 27) with nested 2-fold cross validation (2F-CV) to construct multivariate models that classified participants as male or female. Given our large sample size, using 2-folds minimizes variance and over-fitting while leaving a sufficiently large sample to test model performance. We accounted for age and in-scanner head motion in these models by regressing these covariates from each feature in the training datasets using SurfStat (28) and then applied these model parameters to account for covariates in the testing dataset without data leakage. These models were able to classify participants as male or female with 82.9% accuracy (p<0.0001; **Figure 3a**). Sensitivity and specificity of the model were 0.76 and 0.88, respectively; area under the ROC curve (AUC) was 0.86. To understand which networks contributed the most to the prediction, we summed the positive and negative weights separately across all vertices in each network. This revealed that variation in the functional topography of association networks including the ventral attention network and default mode network contributed the most to the model and were therefore relatively more important in predicting participant sex (**Figure 3B-C**). To determine the importance of a given vertex to the predictive model, we summed the absolute weight across all 17 networks to summarize the prediction weight of each vertex. This summary measure highlighted that regions in association cortex including the temporo-parietal junction, superior parietal lobule, and orbitofrontal cortex were most important in predicting participant sex (**Figure 3D**).

**Figure 3.**
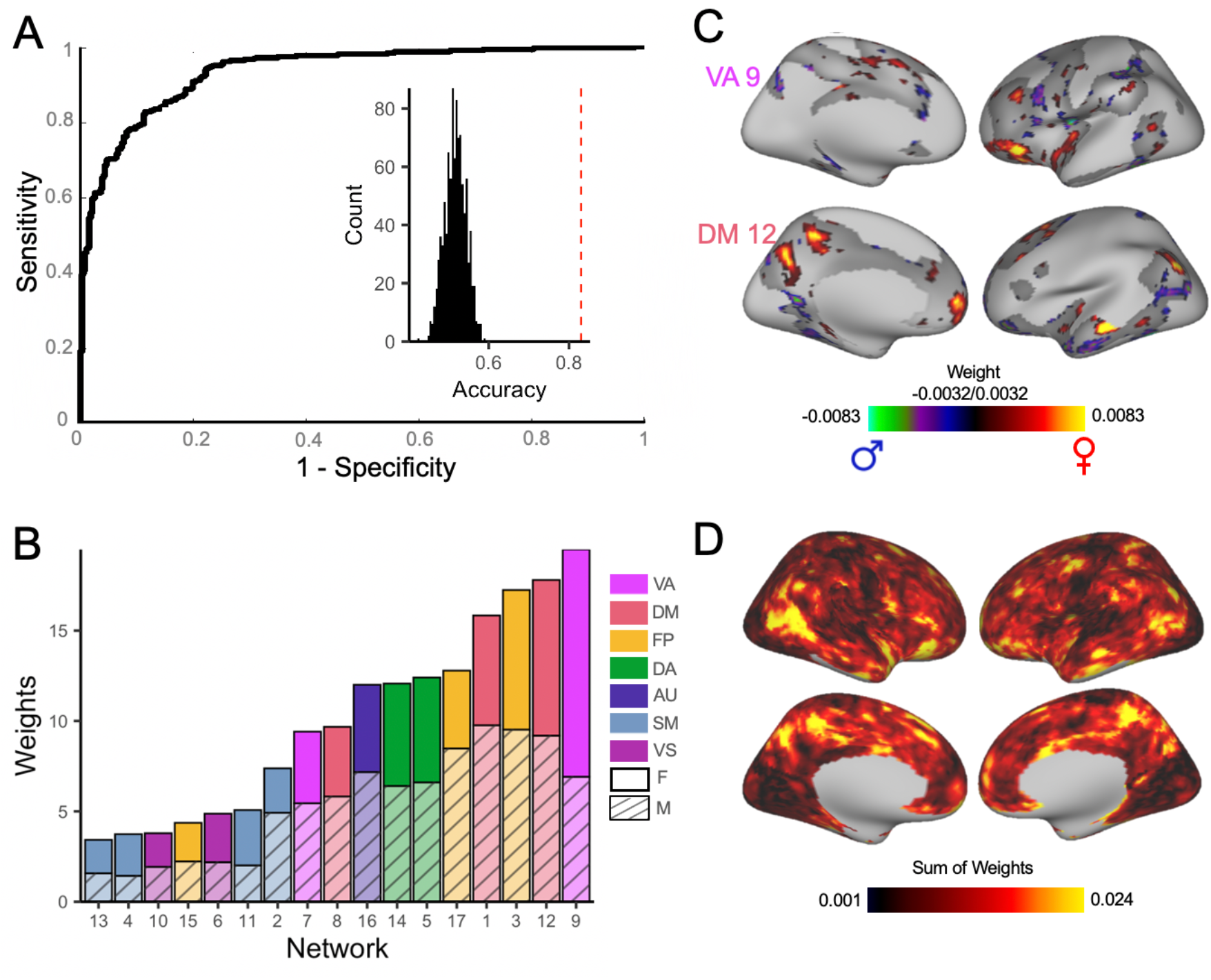
Multivariate pattern analysis using support vector machine predicts subject sex based on functional topography. **A)** Support vector machine (SVM) with nested 2-fold cross validation (2F-CV) was used to construct multivariate models that classified participants as male or female. The ROC curve of the resulting model is depicted. Area under the ROC curve was 0.86. Average sensitivity and specificity of the model were 0.76 and 0.88, respectively. Models classified participants as male or female with 82.9% accuracy. Inset histogram shows distribution of permuted accuracies. Average accuracy is represented by the dashed red line. **B)** To understand which networks contributed the most to the prediction, the positive and negative weights were summed separately across all vertices in each network. The most important topographic features in this model were found in association cortex and were maximal in the ventral attention network and default mode network. **C)** The top 25% of vertices in terms of feature importance in the SVM model are displayed for the ventral attention network and default mode network. **D)** At each location on the cortex, the absolute contribution weight of each network was summed, revealing that association cortex contributed the most to the multivariate model predicting participant sex.

### Mass-univariate analyses yield convergent results

The goal of our multivariate pattern analysis was to classify participant sex using the information contained in all regions jointly. Although multivariate models are ideal for classification problems, their descriptive utility is sometimes limited. The interpretability of features within a multivariate model may be hindered by the inability to determine how the features interact within the model framework due to the high-dimensional nature of the parameter space. In contrast, a traditional mass-univariate analysis describes the relationship between a given factor and brain measures of interest on a regional basis, providing descriptive information complementary to multivariate results. Therefore, we also examined the impact of sex on network topography using a traditional mass-univariate analysis. We used generalized additive models (GAMs) with penalized splines (29) to account for linear and nonlinear developmental effects. We fit a GAM at each vertex to evaluate the impact of sex on network loadings. Age and in-scanner head motion were included as covariates, and age was modeled using a penalized spline. Multiple comparisons were accounted for by controlling the false discovery rate (FDR; *Q*<0.05).

To determine the overall effect of sex at a given vertex, we summed the absolute value of the Z statistic for the effect of sex across all 17 networks. This summary measure highlighted that the impact of sex on topography was greatest in association cortex regions including the temporo-parietal junction, superior parietal lobule, and orbitofrontal cortex (**Figure 4A**). Notably, this result identifying regions where the impact of sex on topography was greatest was convergent with our multivariate pattern analysis, which identified the same regions of association cortex as most heavily weighted in classifying participant sex. We evaluated the significance of the correspondence between this univariate summary measure and the map of summed absolute prediction weights from our machine learning model (**Figure 3D**) using a conservative spin-based spatial randomization test that accounts for spatial autocorrelation (30–33). This analysis revealed that maps of the impact of sex were similar using multivariate and univariate approaches (r=0.86, p_spin_<0.0001; **Figure 4B**). As in our multivariate analysis, the impact of sex using a univariate approach was greatest in association networks including the ventral attention network and default mode network (**Figure 4C-D**). For example, both the SVM and GAMs identified the precuneus as a region with large sex differences in topography; loadings in this region were greater in females for the default mode network, but greater in males for the frontoparietal network (**Figure 4E**). Analyses evaluating the presence of an age-by-sex interaction indicated no significant interactions.

**Figure 4.**
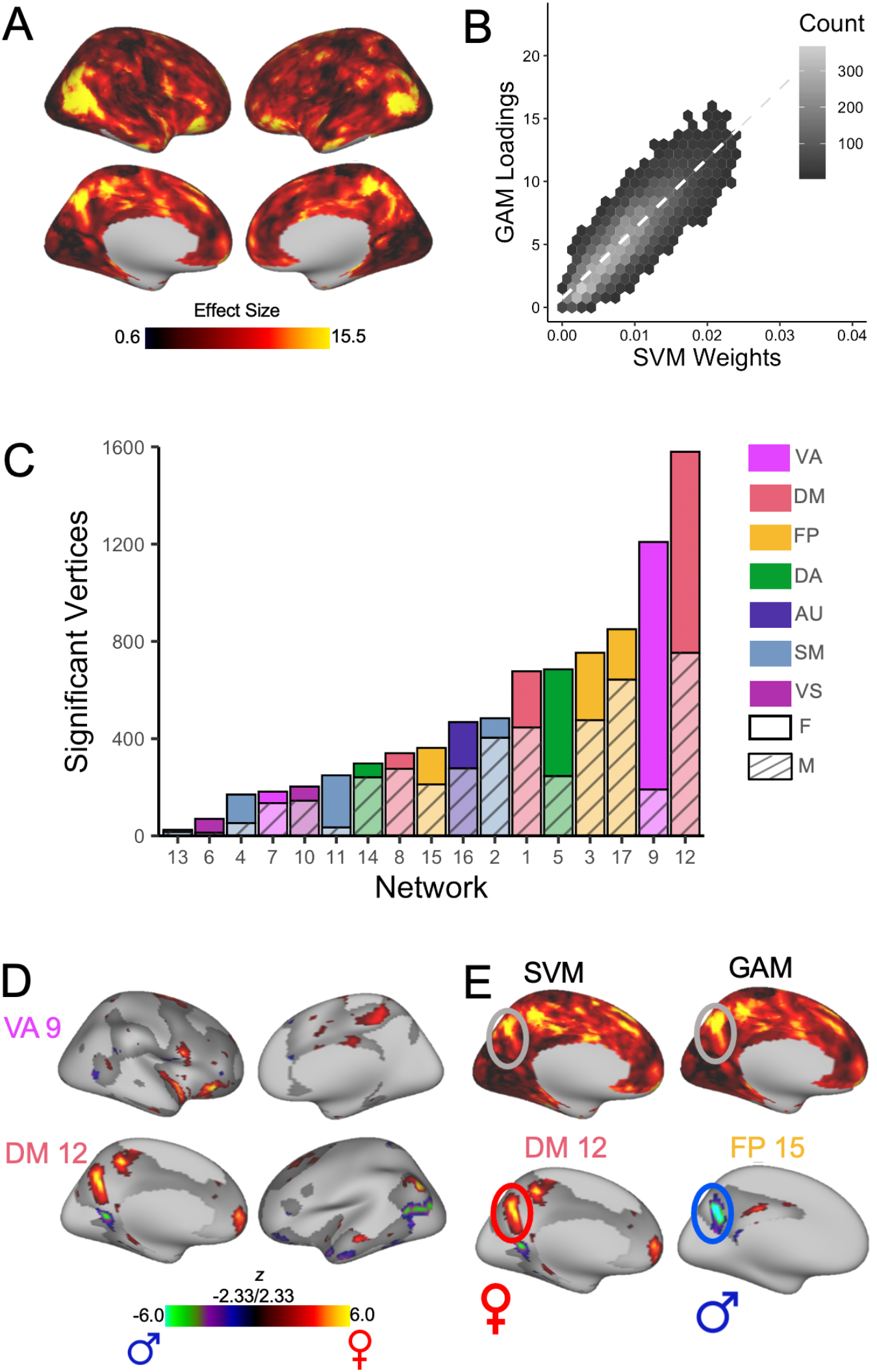
Mass-univariate analyses provide convergent results, identifying significant sex differences in association networks. A generalized additive model (GAM) was fit at each vertex to evaluate the impact of sex on network loadings. Age (modeled using a penalized spline) and motion were included as covariates. Multiple comparisons were accounted by controlling the false discovery rate (Q<0.05). **A)** To determine the overall effect of sex at a given vertex, we summed the absolute sex effect across all 17 networks. This summary measure is depicted and highlights that the impact of sex on topography was greatest in association cortex regions including the temporo-parietal junction, superior parietal lobule, and orbitofrontal cortex. **B)** Hexplot shows agreement between univariate summary measure (GAM loadings in Figure 4A) and multivariate summary measure (SVM weights in Figure 3D) (r=0.86). **C)** To identify networks with the greatest sex differences, the number of vertices in each network with a significant sex effect was summed separately for males and females. This analysis revealed that sex differences were greatest in association networks. **D)** Significant vertices are displayed for the ventral attention network and default mode network, the networks where sex differences were maximal. **E)** Both SVM and GAMs identified the precuneus as a region with large sex differences in topography; loadings in this region were greater for females in the default mode network, but greater in the frontoparietal network for males.

### Gene enrichment analyses link sex differences in topography to X chromosome genes

The above findings indicate that there are robust differences in functional topography between males and females. We next sought to understand the biological basis of these sex differences in topography. Although little is currently known about what factors drive interindividual variation in topography, sex differences in neuroanatomy have previously been attributed to differences in sex chromosome gene expression (22, 23). Therefore, we conducted a chromosomal enrichment analysis to determine if sex differences in topography were spatially coupled to gene expression. We compared the fMRI map of summed absolute prediction weights from our machine learning model to gene expression data from the Allen Human Brain Atlas (across N = 12,986 genes using a 1000-parcel atlas; Methods). We quantified the degree of spatial correspondence using the median rank of a given gene set across the Pearson *r* correlations between each gene’s expression and the fMRI map. As predicted, we observed a significant enrichment of X-chromosome genes (p=0.02; **Figure 5**) - meaning that regions more important in predicting participant sex showed higher correlation with expression of genes on the X-chromosome. This enrichment remained significant in a series of sensitivity analyses that varied the parcellation resolution (p=0.001); that both varied parcellation resolution and used an independent processing pipeline with alternate methods for annotation, filtering, and sample assignment (34–36) (p=0.02); and limited the transcriptomic data to male donors only (p=0.02); see Methods for details.

**Figure 5.**
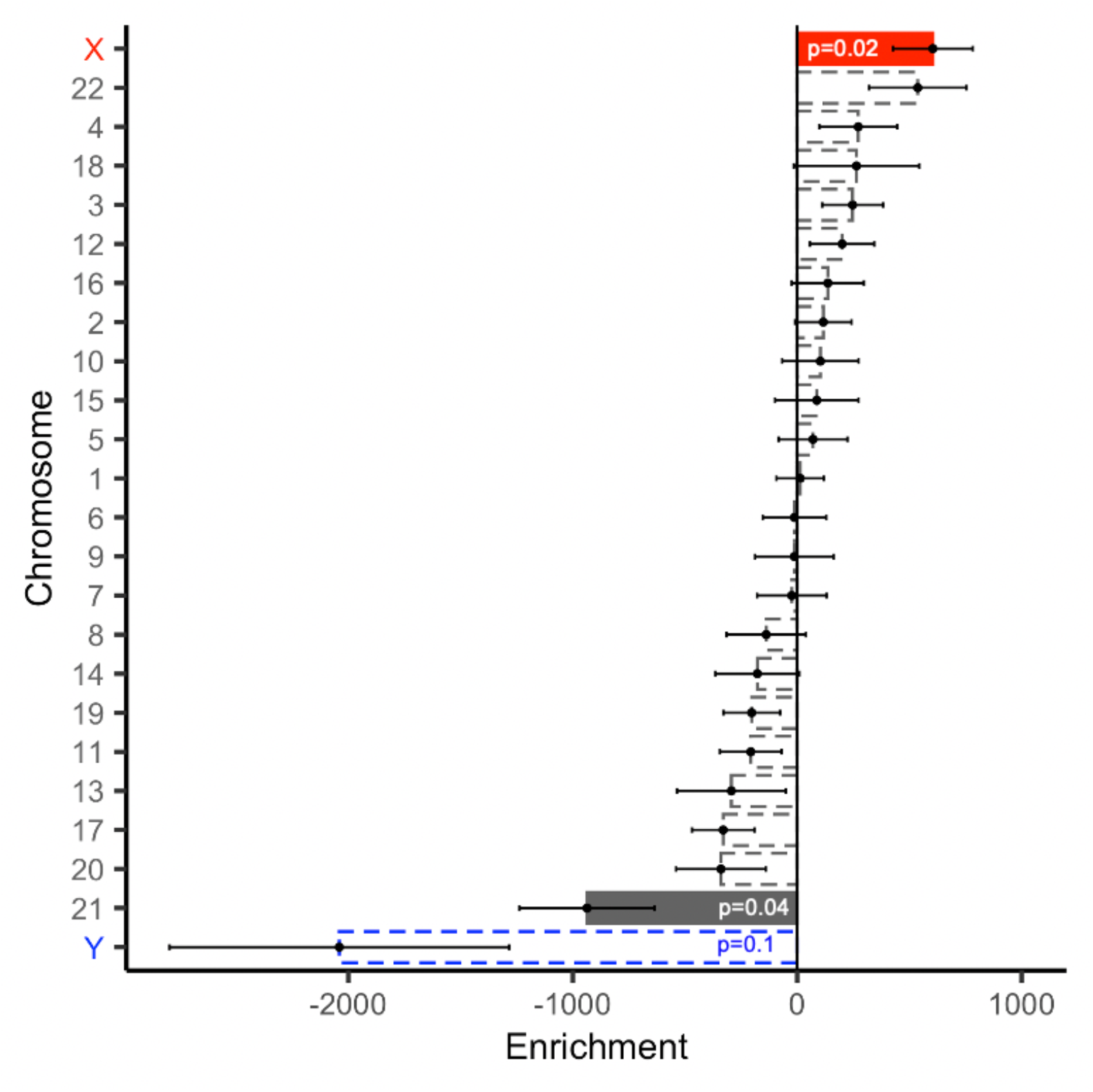
Regions exhibiting sex differences in the multivariate pattern of functional topography are enriched in expression of X-linked genes. To examine the genetic correlates of sex differences in functional topography, we compared the map of summed absolute prediction weights from our machine learning model to gene expression data from the Allen Human Brain Atlas parcellated to the Schaefer 1000 atlas. Point range plot shows the median (point) and standard error (range) rank of each chromosomal gene set. Dashed lines indicate non-significant enrichments. Cortex where sex differences in functional topography were more prominent were significantly correlated with the spatial pattern of expression of genes on the X chromosome.

The above results indicate that sex differences in multivariate patterns of functional topography are correlated with expression of X-linked genes. However, regional differences in cortical gene expression may reflect regional differences in cellular composition of the cortex (37). Therefore, we conducted cell-type specific enrichment analyses to understand the convergent and divergent patterns of discrete underlying gene sets. Using cell-type specific gene sets as assigned in prior work (23), we found that regions more important in classifying participant sex were enriched in expression of astrocytic (p<0.0001) and excitatory neuronal genes (p<0.0001). To obtain a more nuanced understanding of cytoarchitecture, we then assigned cell-types using the finer-grained neuronal sub-class assignments determined by Lake et al. (38). Convergent with the coarser cell type results, regions more important in classifying participant sex were enriched in astrocyte-related genes (p<0.006; **Supplementary Figure 1**) as well as several excitatory neuron sub-classes, including Ex5b (p<0.0001), Ex1 (p<0.0001), Ex3e (p<0.0001), Ex6b (p=0.02), and Ex2 (p=0.03). Notably, these gene sets included numerous X-linked genes (**Supplementary Table 1**). Finally, we conducted a rank-based gene ontology (GO) enrichment analysis using GOrilla (39, 40) to examine functional enrichment. This analysis identified several GO terms relevant to brain anatomy including “neuron part,” “synapse,” and “glutamatergic synapse” (**Supplementary Figure 2**).

## DISCUSSION

In this study, we leveraged machine learning and a large sample of youths to study sex differences in functional network topography. We first demonstrated that sex differences in topography are greatest in association networks, including the ventral attention and default mode networks. Using complex multivariate patterns of functional topography, we were able to predict an unseen participant’s sex with a high degree of accuracy. Chromosomal enrichment analyses revealed that sex differences in multivariate patterns of functional topography were spatially coupled to expression of X-linked genes as well as astrocyte and excitatory neuron cell-type expression signatures. These results identify normative sex differences in the functional topography of association networks and highlight the role of sex as a biological variable in shaping functional brain development in youth.

### Sex differences in functional topography are greatest in association networks

The most robust finding in the present study is that there are significant sex differences in functional topography, and that these differences are greatest in association networks. This finding is in line with prior studies of functional topography that show that inter-individual variation in topography is greatest in association networks (16–20). Importantly, our findings suggest that some significant portion of inter-individual variation in topography is driven by sex. Sex differences in topography were greatest in the default mode network, ventral attention network, and frontoparietal network. Variation in the topography of these networks has been linked to emotional, social, and executive functions (17, 20), all of which are behaviors with documented sex differences (1–3).

To our knowledge, this is the first study to examine sex differences in functional topography. However, our findings are generally convergent with studies examining sex differences in functional connectivity, where standardized network atlases may have aliased differences in topography into measurements of connectivity. The sex differences we found in association cortex topography are in line with several prior studies that have found sex differences in association cortex connectivity (8, 41–45). Specifically, prior studies have documented sex differences in default mode network connectivity (41, 42, 46) and have postulated that sex hormones like estrogen and progesterone might impact default mode connectivity (43, 44, 46, 47). Similarly, our findings of sex differences in the topography of frontoparietal and ventral attention networks align with prior studies that have reported sex differences in functional connectivity of these networks (41, 45) (48). Our multivariate pattern analysis showed that features from the default mode, frontoparietal, and ventral attention networks were most important in classifying participant sex. These results generally cohere with findings from a large study using data from the Human Connectome Project that found that functional connectivity features within the default mode network and frontoparietal network were most important in identifying participant sex (41). Further, abnormal patterns of connectivity involving these association networks have also been associated with mood, fear, and externalizing symptoms (12), dimensions of psychopathology with well documented sex differences. Specifically, abnormalities of default mode, frontoparietal, and ventral attention network connectivity associated with fear symptoms are greater in females (12). Together, these findings suggest that sex differences in topography may contribute to sex differences in cognition and psychopathology, though further work is needed to establish such a relationship.

### Sex differences in topography are associated with X-linked gene expression

The mechanisms by which sex differences in topography arise are likely multifold. Despite the growing interest in inter-individual differences in functional topography, little is currently known about what genetic or environmental factors drive these differences. In prior work, we showed that inter-individual variability in topography aligns with fundamental properties of brain organization including myelin content and cerebral blood flow (20). Here, we built on these findings by linking sex differences in topography to gene expression data. As expected, we found that regions more important in predicting participant sex correlated with expression of genes on the X chromosome. The correspondence between sex differences in topography and the spatial expression pattern of X-linked genes suggests that the observed sex differences in topography are likely in part driven by gene expression. Although to our knowledge no prior studies have examined the genetic basis of sex differences in functional topography, this finding is globally consistent with prior work that has linked sex differences in brain structure to sex chromosome gene expression (22, 23).

Highly ranked X-linked genes included those related to neuron development (*PPEF1* (49), *PCDH19* (50)), neuronal cytoskeletal transport (*DYNLT3* (51)), chloride ion channels (*CLCN5, GABRA3*), metabolism (*GYG2*, *PDK3* (52)), and disease states with neuropsychiatric and cognitive symptoms (*DMD* (53); *DYNLT3* (51), *PCDH19* (50)). Of note, one gene ranking in the top 10 X-linked genes was *PCDH19*, a gene that encodes a protocadherin protein that supports neuronal organization and migration (50). *PCDH19* was also identified as a gene whose spatial expression pattern correlates with sex differences in grey matter volume (22). Mutations in *PCDH19* have been associated with intellectual disability, behavioral problems, and autism spectrum disorder (50). Although speculative, it is plausible that these X-linked genes could influence functional topography through their actions on brain development.

Another mechanism by which sex differences in topography may arise may be via organizational effects of hormones on cytoarchitecture. Our finding that sex differences in topography were spatially coupled to excitatory neuron cell-type gene expression was robust using two separate cell-type categorizations. This result coheres with the extensive rodent and non-human primate literature examining the impact of estradiol on glutamatergic dendritic spine architecture (54) and sex differences in excitatory neurotransmission (55). For example, estrogen is essential to the maintenance of prefrontal cortex dendritic spine density in ovariectomized female rodents and non-human primates (54, 56–58). Estrogen also increases the number of spine synapses in the prefrontal cortex of gonadectomized male rats (59). Similarly, compared to male rodents, female rodents show larger AMPA receptor synaptic responses (60), greater sensitivity to NMDA receptor manipulations (55, 61, 62), and increased expression of NMDA and metabotropic glutamate receptors (55, 63).

We also found that sex differences in topography correlated with astrocytic subtype gene expression. Sex differences in astrocyte structure and astrocytic glutamate release are critical determinants of sex-specific synaptic patterning in sexually dimorphic brain regions, which has been implicated in the male-biased risk for autism spectrum disorder (64). Similarly, androgen receptors play a role in establishing sex differences in astrocyte number and complexity seen in rodents (65). In the context of this literature, we speculate that hormone effects on cytoarchitecture may contribute to sex differences in functional topography.

### Limitations

Certain limitations of the present study should be noted. First, we concatenated three fMRI runs, two of which were task time series where task effects were regressed from the data. Residuals from task-regressed time series, while similar, are not identical to true resting state data and non-linear effects associated with performing the task may therefore not have been removed (66). However, several independent studies have shown that functional networks are primarily defined by individual-specific rather than task-specific factors (67) and that networks present during task performance and at rest are similar (66). Including task-regressed data enabled us to generate individualized networks using 27 min of high-quality data. Time series of this length are necessary to reliably detect individual differences in functional networks (68) and sufficient to create parcellations highly similar to those generated using 380 min of data (69). Second, subcortical and cerebellar networks were not evaluated in this study, as individualized parcellation of these networks requires specialized analysis techniques that are distinct from those applied to the cortex (70, 71), rendering comparative or conjunctive analyses with NMF difficult to perform and interpret. Future work should evaluate sex differences in topography in subcortical and cerebellar networks, which are critical for behaviors with known sex differences including emotional regulation and executive function.

Third, correspondence between sex differences in topography and gene expression were assessed at the group rather than individual level, though evaluating this relationship on a within-subject basis in a large, developmental sample is precluded by the necessity of postmortem samples for gene expression profiling. Fourth, using the Allen Human Brain Atlas introduces several inherent limitations including the use of microarray to quantify gene expression, asymmetric sampling, small sample size, donor age, and most notably, donor sex. The Allen includes postmortem samples from five male donors and one female donor. However, our findings regarding the correspondence between sex differences in topography and X-linked gene expression were robust to sensitivity analyses leaving out the female donor, and prior studies have similarly leveraged the AHBA to examine sex differences in independent neuroimaging samples (22, 23). Nevertheless, replication of these findings in a sex-balanced sample will be important when such spatially comprehensive maps of cortical gene expression are available.

Fifth, sex was assessed using a binary self-report question, and we therefore did not have a sufficient sample to examine functional topography of intersex youths. Furthermore, it should be noted that existing data and theory suggest that binary sex classification may not be useful and that brains are complex mosaics of male and female characteristics (72).

### Conclusions

In summary, we identified normative developmental sex differences in the functional topography of personalized association networks. These results suggest that inter-individual variation in functional topography is in part driven by sex. Further, our findings suggest that sex differences in topography are likely in part linked to gene expression. Future work should examine if the sex differences in the topography of personalized networks explain normative variation in socioemotional or cognitive functions. The relationship between topography and sex differences in psychopathology is also a clear area for future research, as this may identify sexspecific biomarkers of risk for psychiatric disorders.

## MATERIALS AND METHODS

Methods regarding sample construction, image acquisition and processing, and individualized network parcellation were as previously described (20) and are summarized briefly below.

### Participants

In this report, we considered the entire cross-sectional sample of 1,601 subjects imaged as part of the Philadelphia Neurodevelopmental Cohort (21). From this sample, 340 subjects were excluded due to clinical factors, including medical disorders that could affect brain function, current use of psychoactive medications, prior inpatient psychiatric treatment, or an incidentally encountered structural brain abnormality. An additional 568 subjects were excluded because of low quality or missing structural, resting-state, n-back, or emotion identification imaging data; a functional run was excluded if mean relative root mean square (RMS) framewise displacement was higher than 0.2mm, or it had more than 20 frames with motion exceeding 0.25 mm (73, 74). The final sample included in the analyses comprised 693 participants of which 301 were male and 392 were female. This sample of participants and their individualized networks are the same as those included in our prior report on individual variation in functional network topography (20). All subjects or their parent/guardian provided informed consent, and minors provided assent. All study procedures were approved by the Institutional Review Boards of both the University of Pennsylvania and the Children’s Hospital of Philadelphia.

### Image acquisition

As previously described (21), all MRI scans were acquired using the same 3T Siemens Tim Trio whole-body scanner and 32-channel head coil at the Hospital of the University of Pennsylvania.

#### Structural MRI

Prior to functional MRI acquisitions, a 5-min magnetization-prepared, rapid acquisition gradient-echo T1-weighted (MPRAGE) image (TR =1810ms; TE=3.51ms; TI=1100ms, FOV=180 x 240mm^2^, matrix=192 x 256, effective voxel resolution=0.9 x 0.9 x 1mm^3^) was acquired.

#### Functional MRI

We used one resting-state and two task-based (i.e., *n*-back and emotion recognition) fMRI scans as part of this study. All fMRI scans were acquired with the same single-shot, interleaved multi-slice, gradient-echo, echo planar imaging (GE-EPI) sequence sensitive to BOLD contrast with the following parameters: TR = 3000 ms; TE = 32 ms; flip angle = 90°; FOV = 192 x 192 mm^2^; matrix = 64 x 64; 46 slices; slice thickness/gap = 3/0 mm, effective voxel resolution = 3.0 x 3.0 x 3.0 mm^3^. Resting-state scans had 124 volumes, while the *n*-back and emotion recognition scans had 231 and 210 volumes, respectively. Further details regarding the *n*-back (75) and emotion recognition (76) tasks have been described in prior publications.

#### Field map

In addition, a B0 field map was derived for application of distortion correction procedures, using a double-echo, gradient-recalled echo (GRE) sequence: TR = 1000ms; TE1 = 2.69 ms; TE2 = 5.27 ms; 44 slices; slice thickness/gap = 4/0 mm; FOV = 240 mm; effective voxel resolution = 3.8 x 3.8 x 4 mm.

### Image processing

The structural images were processed using FreeSurfer (version 5.3) to allow for the projection of functional time series to the cortical surface (77). Functional images were processed using a top-performing preprocessing pipeline implemented using the eXtensible Connectivity Pipeline (XCP) Engine (73), which includes tools from FSL (78, 79) and AFNI (80). This pipeline included 1) correction for distortions induced by magnetic field inhomogeneity using FSL’s FUGUE utility, 2) removal of the initial volumes of each acquisition (i.e., 4 volumes for resting-state fMRI and 6 volumes for emotion recognition task fMRI), 3) realignment of all volumes to a selected reference volume using FSL’s MCFLIRT, 4) interpolation of intensity outliers in each voxel’s time series using AFNI’s 3dDespike utility, 5) demeaning and removal of any linear or quadratic trends, and 6) co-registration of functional data to the high-resolution structural image using boundary-based registration. Images were de-noised using a 36-parameter confound regression model that has been shown to minimize associations with motion artifact while retaining signals of interest in distinct sub-networks. This model included the six framewise estimates of motion, the mean signal extracted from eroded white matter and cerebrospinal fluid compartments, the mean signal extracted from the entire brain, the derivatives of each of these nine parameters, and quadratic terms of each of the nine parameters and their derivatives. Both the BOLD-weighted time series and the artifactual model time series were temporally filtered using a first-order Butterworth filter with a passband between 0.01 and 0.08 Hz to avoid mismatch in the temporal domain (81). Furthermore, to derive ‘‘pseudo-resting state’’ time series that were comparable across runs, the task activation model was regressed from n-back or emotion recognition fMRI data (66). The task activation model and nuisance matrix were regressed out using AFNI’s 3dTproject.

For each modality, the fMRI time series of each individual were projected to each subject’s FreeSurfer surface reconstruction and smoothed on the surface with a 6-mm full-width half-maximum (FWHM) kernel. The smoothed data was projected to the *fsaverage5* template, which has 10,242 vertices on each hemisphere (18,715 vertices in total after removing the medial wall). Finally, we concatenated the three fMRI acquisitions, yielding time series of 27 minutes, 45 seconds (555 time points) in total.

As in prior work, we removed vertices with low signal-to-noise ratio (SNR) (31, 82, 83). To calculate a whole-brain SNR map, we extracted the first frame of acquisition (post-steady state magnetization) of each of the three runs for all participants. Next, we normalized each image to a mode of 1,000 and then averaged all of these images; this average image was further normalized to a mode of 1,000. A mean BOLD signal of 800 or less represents a substantial attenuation of signal (83), applying this threshold resulted in the exclusion of low SNR locations, which were primarily located in orbitofrontal cortex and ventral temporal cortex. Within this mask, 17,734 vertices were included in subsequent analyses.

### Regularized non-negative matrix factorization

As previously described in detail (18), we used non-negative matrix factorization (NMF) (24) to derive individualized functional networks. NMF factors the data by positively weighting cortical elements that covary to yield a highly specific and reproducible parts-based representation (24, 32). Our approach was enhanced by a group sparsity consensus regularization term that preserves inter-individual correspondence, as well as a data locality regularization term that makes the decomposition robust to imaging noise, improves spatial smoothness, and enhances functional coherence of subject-specific functional networks (see Li et al., 2017 (18) for method details; see also: https://github.com/hmlicas/Collaborative_Brain_Decomposition). Because NMF requires inputs to be nonnegative values, we re-scaled the data by shifting time series of each vertex linearly to ensure all values were positive (18). To avoid features in greater numeric ranges dominating those in smaller numeric range, we further normalized the time series by its maximum value so that all the time points have values in the range of [0, 1].

Given a group of *n* subjects, each having fMRI data *X^i^* ∈ *R*^×^, *i* = 1,…, *n*, consisting of *S* vertices and *T* time points, we aimed to find *K* non-negative functional networks 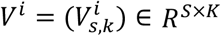 and their corresponding time series 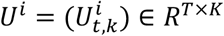 for each subject, such that

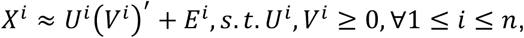

where (*V^i^*) is the transpose of (*V^i^*), and *E^i^* is independently and identically distributed (i.i.d) residual noise following Gaussian distribution with a probability density function of 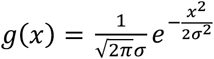. Both *U^i^* and *V^i^* were constrained to be non-negative so that each functional network does not contain any anti-correlated functional units (24). A group consensus regularization term was applied to ensure inter-individual correspondence, which was implemented as a scale-invariant group sparsity term on each column of *V^i^* and formulated as

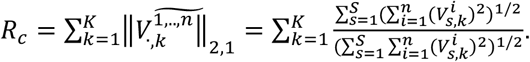

The data locality regularization term was applied to encourage spatial smoothness and coherence of the functional networks using graph regularization techniques (84). The data locality regularization term was formulated as

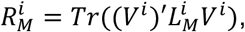

where 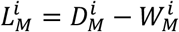 is a Laplacian matrix for subject *i*, 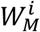 is a pairwise affinity matrix to measure spatial closeness or functional similarity between different vertices, and 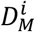 is its corresponding degree matrix. The affinity between each pair of spatially connected vertices (i.e., vertices *a* and *b*) was calculated as 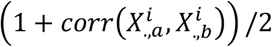, where 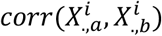 is the Pearson correlation coefficient between time series 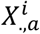 and 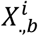, and others were set to zero so that 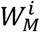 has a sparse structure. We identified subject specific functional networks by optimizing a joint model with integrated data fitting and regularization terms formulated by

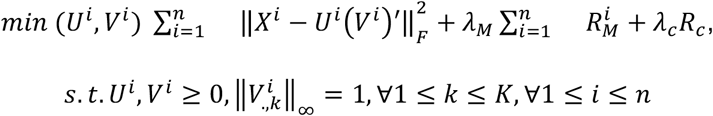

where 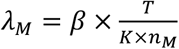 and 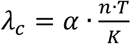 are used to balance the data fitting, data locality, and group consensus regularization terms, *n_M_* is the number of neighboring vertices, *α* and *β* are free parameters. Here, we used the same parameter settings as those used in prior validation studies (18).

### Defining individualized networks

Our approach for defining individualized networks included three steps (**Figure 1**). The first two steps yielded a consensus group atlas. In the third step, this group atlas was used to define individualized networks for each participant. The whole-brain was decomposed into 17 networks for correspondence with commonly used atlases and prior work (17, 20, 25, 26).

#### Step 1: Group network initialization

Although individuals exhibit distinct network topography, they are also broadly consistent (19, 67). Therefore, we first generated a group atlas and used this group atlas as an initialization for individualized network definition. By doing so, we ensured spatial correspondence across all subjects. This strategy has been applied in prior work for individualized network definition (17, 20, 25). To avoid the group atlas being driven by outliers and to reduce the computational memory cost, a bootstrap strategy was utilized to perform the group-level decomposition multiple times on a subset of randomly selected subjects. Subsequently, the resulting decomposition results were fused to obtain one robust initialization that is highly reproducible. As in prior work (18), we randomly selected 100 subjects and temporally concatenated their time series, resulting in a time series matrix with 55,500 rows (time-points) and 17,734 columns (vertices). The choice of sub-sample size did not impact results (sub-samples of 100, 200, and 300 were previously evaluated (20)). We applied the above-mentioned regularized non-negative matrix factorization method with a random non-negative initialization to decompose this matrix (24). A group-level network loading matrix *V* was acquired, which had 17 rows and 17,734 columns. Each row of this matrix represents a functional network, while each column represents the loadings of a given cortical vertex. This procedure was repeated 50 times, each time with a different subset of subjects (18); this yielded 50 different group atlases.

#### Step 2: Group network consensus

Next, we used spectral clustering to combine the 50 group network atlases into one robust and highly reproducible group network atlas (18). Specifically, we concatenated the 50 group parcellations together across networks and acquired a matrix with 850 rows (i.e., functional networks, abbreviated as FN) and 17,734 columns (i.e., vertices). Inter-network similarity was calculated as

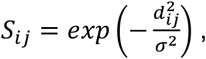

where *d_ij_* = 1 – *corr*(*FN_i_, FN_j_*), *corr*(*FN_i_, FN_j_*) is Pearson correlation coefficient between *FN_i_* and *FN_j_*, and *σ* is the median of *d_ij_* across all possible pairs of FNs. Then, we applied normalized-cuts (84) based spectral clustering method to group the 850 FNs into 17 clusters. For each cluster, the FN with the highest overall similarity with all other FNs within the same cluster was selected as the most representative. The final group network atlas was composed of the representatives of these 17 clusters.

#### Step 3: Individualized networks

In this final step, we derived each individual’s specific network atlas using regularized NMF based on the acquired group networks (17 x 17,734 loading matrix) as initialization and each individual’s specific fMRI times series (555 x 17,734 matrix). See Li et al., 2017 (18) for optimization details. This procedure yielded a loading matrix V (17 x 17,734 matrix) for each participant, where each row is a functional network, each column is a vertex, and the value quantifies the extent each vertex belongs to each network. This probabilistic (soft) definition can be converted into discrete (hard) network definitions for display by labeling each vertex according to its highest loading.

### Multivariate Pattern Analysis

We used a linear support vector machine (LSVM) as implemented in LIBSVM (85) to construct multivariate models that classified participants as male or female. A free parameter *C* determines the balance between the training errors and the generalizability of the LSVM classification model. We evaluated the classification using a nested 2-fold cross validation (2F-CV), with the inner 2F-CV determining the optimal parameter *C* for the SVM classifier and the outer 2F-CV estimating the generalizability of the model. Given the large sample size in this study, using 2-folds minimizes variance and over-fitting while leaving a sufficiently large sample to test model performance.

In the outer 2F-CV, the data was randomly divided into 2 subsets. We initially used subset 1 as the training set, with subset 2 used as the testing set. We accounted for age and in-scanner head motion by regressing these effects from each feature in the training datasets using SurfStat (28) and then applied the acquired coefficients to regress the effects in testing dataset. Each feature was linearly scaled between zero and one across the training dataset; these scaling parameters were then applied to scale the testing dataset (86, 87).

Within each outer 2F-CV loop, we applied inner 2F-CV loops to determine the optimal *C*. To do this, the training set of the outer 2F-CV loop was again randomly divided into 2 subsets; one subset was selected to train the model with a given *C* in the range [2^-5^, 2^-4^,…, 2^9^, 2^10^] (i.e., 16 values in total) (88), and the remaining subset was used to test the model. This procedure was repeated 2 times such that each subset was used once as the testing subset, resulting in two inner cross validations in total. The accuracies were calculated for each *C* value and then averaged across the two inner cross validations. The *C* with the highest inner prediction accuracy was chosen as the optimal *C* (86, 89). Then, we trained a model using the optimal *C* and all participants in the training set (subset 1), and then used that model to predict the sex of all participants in the testing set (subset 2).

We repeated the above procedure using subset 2 as the training set and subset 1 as the testing set. Because the split between training and testing sets was chosen randomly, the nested 2-fold cross validation was performed 100 times to reduce the impact of group assignment. The results of these 100 nested cross validations were then averaged. This procedure yielded an overall classification accuracy score for classifying males or females on the basis of their functional topography. Each SVM also produced a vector of feature weights for each functional network that described how heavily weighted a given topographic feature was within the multivariate model.

#### Significance of prediction performance

Permutation testing was used to evaluate if the prediction performance was significantly better than expected by chance (90). The predictive framework was repeated 1,000 times. In each run, we permuted sex across the training subset without replacement. Significance was determined by ranking the actual prediction accuracy (the average across 100 runs) versus the permuted distribution; the *p-value* was the proportion of permutations that showed a higher value than the actual accuracy value for the real data.

#### Interpreting model feature weights

The nested 2F-CV was repeated 100 times, yielding 200 weight maps. Averaging these 200 weight maps, the features with a nonzero mean weight can be understood as contributing features in the prediction model (86, 90), with the absolute value of the weight quantifying the contribution of the features to the model (90). To understand which networks contributed the most to the prediction, we summed the positive and negative weights separately across all vertices in each network. As each vertex had 17 loading values (one for each network), we summed the absolute weight across all 17 networks to summarize the prediction weight of each vertex. This sum represents the importance of a given vertex to the predictive model.

### Univariate associations of network topography with sex

The goal of a multivariate pattern analysis is to predict an outcome using the information contained in all regions jointly. In contrast, the goal of a traditional mass-univariate analysis is to describe the relationship between a given factor and brain measures of interest on a regional basis. Although multivariate models are ideal for classification problems, their descriptive utility is limited. While it is possible to identify the most heavily weighted features within a model, it is not possible to directly visualize their action within the model framework due to the highdimensional nature of the parameter space. As such, we used both multivariate (predictive) and mass-univariate (descriptive) approaches, which are complementary and provide converging evidence.

We evaluated mass-univariate associations between sex and network topography using generalized additive models (GAMs) with penalized splines (29) to account for linear and nonlinear developmental effects. GAMs estimate nonlinearities using restricted maximum likelihood (REML), penalizing nonlinearity in order to avoid over-fitting the data. We fit a GAM at each vertex to evaluate the impact of sex on network loadings. Age and in-scanner head motion were included as covariates, and age was modeled using a penalized spline. As we considered three functional runs, in-scanner motion was summarized as the grand mean of the mean relative RMS displacement of each functional run. Multiple comparisons were accounted for by controlling the false discovery rate (FDR; *Q*<0.05).

### Spatial randomization testing (spin test)

To evaluate the significance of the correspondence between our multivariate and univariate results, we compared a map of summed absolute prediction weights from our machine learning model (**Figure 4D**) to a map of summed effect sizes from the GAMs (**Figure 5A**). We compared these maps using the spin test (30–33) (https://github.com/spin-test/spin-test). The spin test is a spatial randomization method based on angular permutations of spherical projections at the cortical surface. The spin test generates a null distribution of randomly rotated brain maps that preserve spatial features and the spatial covariance structure of the original map. This procedure is therefore far more conservative than randomly shuffling locations, which destroys the spatial covariance structure of the data and produces an unrealistically weak null distribution. The spin-based *p*-value was calculated as the proportion of times that the observed correlation was higher than the null distribution of correlation values from rotated parcellations.

### Gene enrichment analysis

To examine the transcriptomic correlates of sex differences in functional topography, we compared the map of summed absolute prediction weights from our machine learning model to gene expression data from the Allen Human Brain Atlas (91). Publicly available microarray gene expression data processed in line with recent benchmarking recommendations and parcellated to the Schaefer1000 atlas were downloaded from https://figshare.com/articles/dataset/AHBAdata/6852911. Details regarding gene processing including gene information re-annotation, data filtering, probe selection, sample assignment, data normalization, and gene filtering are described in Arnatkevičiuté *et al*. (92). Of the available parcellations of processed Allen data, the Schaefer1000 parcellation was selected for primary comparison with topography given the granularity of topographic features. As only two of the six donor brains were sampled from both hemispheres, analyses were restricted to the left hemisphere to minimize variability of samples available across regions (92).

#### Chromosomal enrichment analysis

We used ranked gene lists to test if the spatial expression pattern of a given gene set was nonrandomly related to the spatial pattern of sex differences in functional topography. As in prior studies (22, 23, 36), we quantified the degree of spatial correspondence using the median gene set rank. Genes were assigned to chromosomes as annotated in Richiardi *et al*. (93). We calculated the median ranks for 24 non-overlapping gene sets: each autosome (chromosomes 1 through 22), chromosome X, and chromosome Y. For each chromosomal gene set, we compared the observed median rank to a null distribution of median ranks calculated from 1,000 same-sized scrambled lists generated by randomly reordering the original ranked list. The *p-value* from this non-parametric permutation test was the proportion of permutations with a more extreme value than the median rank of the real data.

To test the robustness of our chromosomal enrichments, we conducted a series of sensitivity analyses including varying the processing strategy and parcellation resolution, as well as leaving the female donor out. Specifically, we replicated our results using publicly available gene expression data processed by Arnatkevičiuté *et al*. (92) and parcellated to the Schaefer300 atlas. For consistency with prior work, we also parcellated Allen data to the Scaefer400 atlas using an independent processing pipeline (34–36) and computed gene expression matrices with and without the female donor.

#### Cell-type specific expression analyses and gene ontology

Because regional differences in cortical gene expression may reflect regional differences in cellular composition of the cortex (37), we conducted cell-type specific enrichment analyses to understand the convergent and divergent patterns of discrete underlying gene sets. As in chromosomal enrichments, we used ranked gene lists to test if the spatial expression pattern of a cell-type specific gene set was nonrandomly related to the spatial pattern of sex differences in functional topography. Non-parametric permutation testing assessed the significance of median ranks of cell-type specific gene sets. Gene sets for each cell-type were first assigned according to categorizations determined by Seidlitz *et al*. (23). To obtain a more nuanced understanding of cytoarchitecture, we then used the finer-grained neuronal sub-class assignments determined by Lake *et al*. (38). In both cases, only brain expressed genes (23) were considered, defined by expression levels in the Human Protein Atlas (94, 95).

Finally, we also conducted a rank-based gene ontology (GO) enrichment analysis using GOrilla (39, 40) to examine functional enrichment.

### Visualization

Connectome Workbench (version: 1.3.2) provided by the human connectome project (https://www.humanconnectome.org/software/connectome-workbench; (96)) was used to visualize the brain surface.

### Data & code availability

The PNC data is publicly available in the Database of Genotypes and Phenotypes: accession number: phs000607.v3.p2; https://www.ncbi.nlm.nih.gov/projects/gap/cgi-bin/study.cgi?study_id=phs000607.v3.p2. All analysis code is available here: https://sheilashanmugan.github.io/funcParcelSexDiff1/.

## Acknowledgements & Funding Sources

This study was supported by R01MH120482, R01MH113550, R01EB022573, R01MH107703, RF1MH116920, R01MH112847, P50MH096891, R01MH11186, K01MH102609, R01MH107235, R01MH112070 R01NS085211, RC2MH08998, RC2MH089924, R25MH119043, K08MH120564, T32MH019112 and the Penn/ CHOP Lifespan Brain Institute.

## Supplementary Tables

**Supplementary Table 1.** Cell type sub-classes with expression patterns that correlate with sex differences in topography contain several X-linked genes. Regions exhibiting sex differences in the multivariate pattern of functional topography are enriched in astrocyte-related genes, as well as several excitatory neuron sub-classes, including Ex5b, Ex1, Ex3e, Ex6b, and Ex2. The X-linked genes contained in each of these sub-classes is listed in the table.

| Sub-class | X-linked genes (N) | X-linked genes |
| --- | --- | --- |
| Ast | 2 | *GPM6B, OPHN1* |
| Ex1 | 2 | *DIAPH2, GRIA3* |
| Ex2 | 0 | - |
| Ex3e | 16 | *ATP6AP1, ATP6AP2, BEX1, BEX2, BEX4, CDKL5, GDI1, MAP7D2, MORF4L2, NDUFA1, SYN1, SYP, TMSB4X, TSPAN7, TSPYL2, USP11* |
| Ex5b | 2 | *DMD, GRIA3* |
| Ex6b | 3 | *DMD, GRIA3, PCDH11X* |

## Supplementary Figures

**Supplementary Figure 1.**
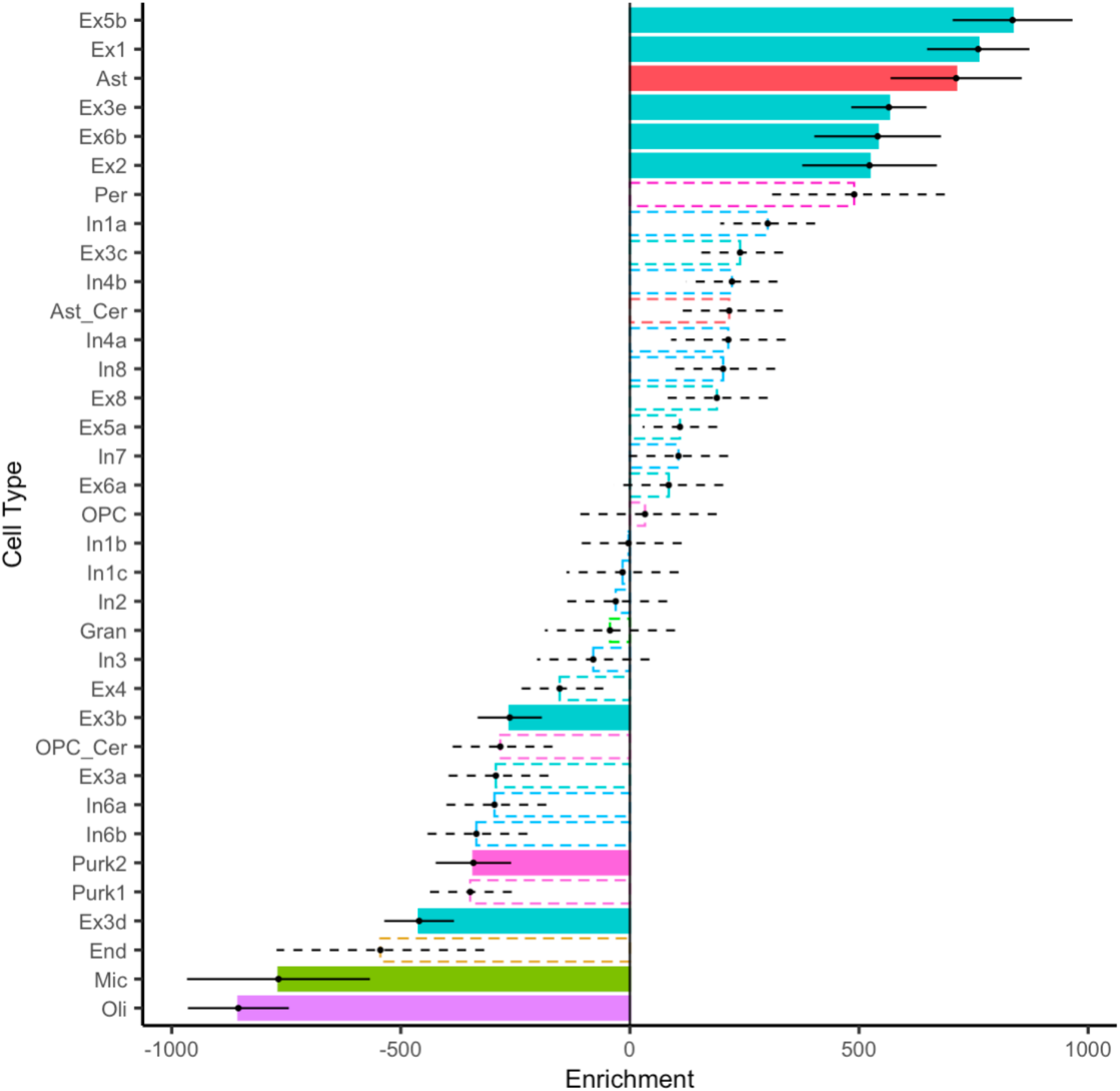
Regions exhibiting sex differences in the multivariate pattern of functional topography are enriched in expression of excitatory neuronal and astrocytic related genes. We conducted cell-type specific enrichment analyses to understand the convergent and divergent patterns of discrete underlying gene sets. We compared the map of summed absolute prediction weights from our machine learning model to gene expression data from the Allen Human Brain Atlas parcellated to the Schaefer 1000 atlas. We assigned celltypes using the neuronal sub-class assignments determined by Lake et al. Point range plot shows the median (point) and standard error (range) rank of each cell type gene set. Dashed lines indicate non-significant enrichments. Regions more important in classifying participant sex were enriched in astrocyte-related genes and several excitatory neuron related gene sets including Ex5b, Ex1, Ex3e, Ex6b, and Ex2. Ast= astrocyte, Ast_cer= cerebellar-specific astrocytes, End= endothelial cells, Ex= excitatory neuron, Gran= cerebellar granule cells, In= inhibitory neuron, Mic= microglia, Oli= oligodendrocytes, OPC= oligodendrocyte progenitor cells, OPC_cer= cerebellar-specific oligodendrocyte progenitor cells, Per= pericytes, Purk= cerebellar Purkinje cells.

**Supplementary Figure 2.**
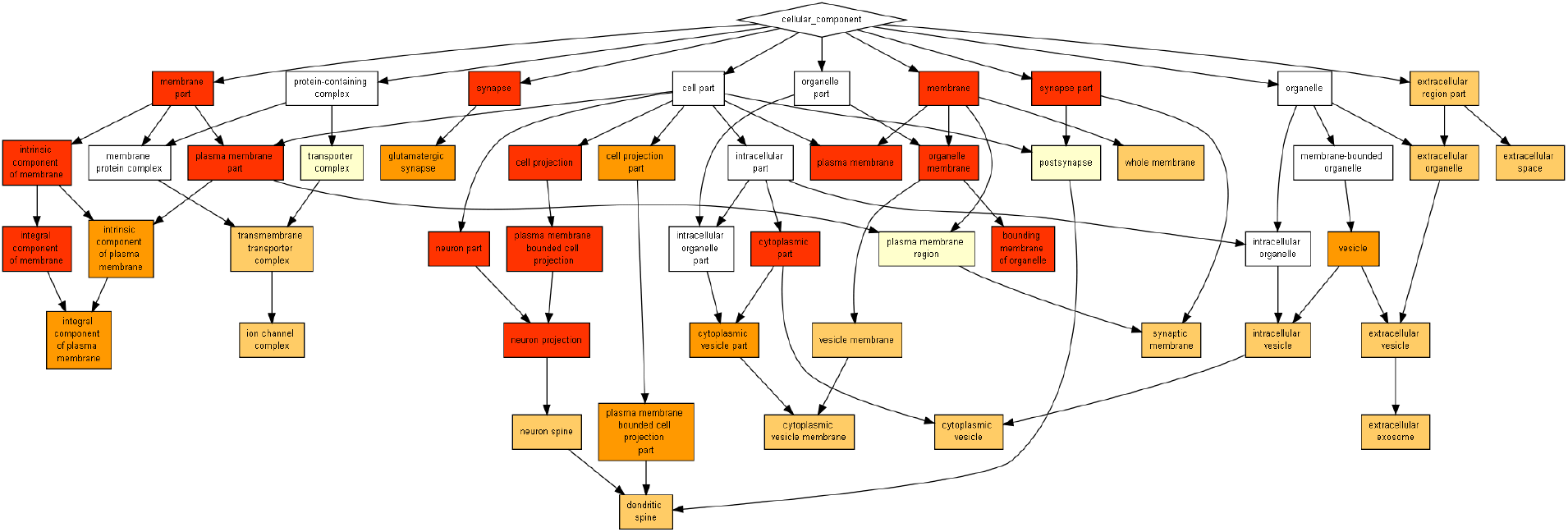
Regions exhibiting sex differences in the multivariate pattern of functional topography are enriched in expression of neuron related genes. We explored whether areas with prominent sex differences in topography show enriched annotation for specific biological processes, cellular components, and molecular functions. We conducted a rank-based gene ontology (GO) enrichment analysis using GOrilla to examine functional enrichment. Full output of cellular component GO terms is depicted. This analysis identified several GO terms relevant to brain anatomy including “neuron part,” “synapse,” and “glutamatergic synapse.”

